# Oxidative Stress-Induced Immunogenic Cell Death Enhances Whole-Cell Vaccine Efficacy in a Syngeneic Pancreatic Cancer Model

**DOI:** 10.64898/2026.02.09.704828

**Authors:** Maryam Katoueezadeh, Yazhini Thinakaran, Mahdi Hejazi Laein, Ranjith Iyappan, SoFong Cam Ngan, Jake Baker, Ridhi Patel, Pazhanichamy Kalailingam, Rebecca E. K. Macpherson, Panagiota Klentrou, Evangelia Litsa Tsiani, Jee Keem Low, Neil E. McCarthy, Siu Kwan Sze

## Abstract

Pancreatic ductal adenocarcinoma (PDAC) is a highly aggressive cancer, with limited therapeutic options and extremely high mortality rates. While immune checkpoint blockade (ICB) therapy is effective in many types of human cancers, responses in PDAC patients remain poor, partly due to the weak immunogenicity of PDAC tumors. We hypothesized that a whole-cell PDAC vaccine could improve anti-tumor responses if optimized to expose a more stimulatory repertoire of tumor antigens. To test this, we used murine Panc02 pancreatic cancer cells to screen several stress-inducing treatments (UV, hypoxia, heat shock, and hydrogen peroxide [H_2_O_2_]), among which low-dose oxidative stress (0.05% H_2_O_2_ for 2h) was identified as the optimal inducer of immunogenic cell death (including increased surface calreticulin, ERp57 exposure, HMGB1 release and MHC class I expression). We then prepared a whole-cell vaccine of fixed H_2_O_2_-treated Panc02 cells, which induced robust tumor-specific immunity in C57BL/6 mice bearing syngeneic Panc02 tumors. Vaccine-treated mice displayed a significant increase in tumor-reactive IFNγ+ T cells, as well as extensive tumor infiltration by CD4 + and CD8 + T cells and NCR1^+^ NK cells. When used prophylactically, the vaccine significantly delayed tumor growth and extended survival, whereas therapeutic application markedly slowed tumor progression. Importantly, combining the whole-cell Panc02 vaccine with anti-PD-1 therapy induced complete tumor regression in a subset of animals. Together, these data demonstrate that controlled oxidative stress can convert autologous tumor cells into an effective whole-cell vaccine without the need for genetic modification or prior neoantigen identification, offering a scalable strategy for personalized immunotherapy in PDAC.

**STATEMENT OF SIGNIFICANCE:** This study demonstrated that oxidative stress-induced immunogenic cell death reprograms pancreatic tumor cells to induce danger signaling and enhance antigen presentation, thereby promoting immune infiltration and sensitizing tumors to PD-1 blockade.

## INTRODUCTION

Pancreatic ductal adenocarcinoma (PDAC) is one of the deadliest forms of cancer with the poorest 5-year survival rate of less than 13% among all cancer types[1]. Late diagnosis and aggressive clinical course of PDAC contribute to poor outcomes, but even localized PDAC often recurs rapidly after surgery[2]. Treatment options are limited, partly due to PDAC being an immunologically ‘cold’ tumor, in which the microenvironment is dominated by suppressive cell types and dense stroma[3]. As a result, potent immunotherapies that have revolutionized the treatment of other cancer types have largely failed for PDAC. Clinical trials of immune checkpoint blockade (ICB) targeting PD-1/PD-L1 or CTLA-4 have shown minimal benefits in PDAC patients, with no significant improvement in survival when added to chemotherapy[4]. This is thought to be a consequence of poor T-cell infiltration and antigen presentation in PDAC lesions [5] hence, new strategies are needed to convert these malignancies into immunogenically ‘hot’ tumors.

One approach is to use cancer vaccines that can prime and expand tumor-specific T cells *in vivo*[6]. In this context, whole-cell-based methods can potentially expose the full repertoire of PDAC tumor antigens and patient-specific neoantigens for immune targeting[7]. A key precedent for this concept is the GVAX platform, which uses irradiated allogeneic PDAC cells that are engineered to secrete immunostimulatory GM-CSF[8]. In early clinical studies, a single dose of GVAX was safe and sufficient to recruit previously absent lymphocytes into PDAC lesions, effectively converting ‘cold’ tumors into immunologically active sites[9]. Combining GVAX with immune modulators has since yielded encouraging results: a Phase 1b trial of GVAX treatment together with the CTLA-4 inhibitor ipilimumab improved patient survival compared to ipilimumab alone[10]. More recent trials incorporating PD-1 blockade and CD137 agonists with GVAX therapy have led to enhanced activation of intratumoral T cells[11]. These findings demonstrate that whole-cell vaccines can partially overcome immune resistance in PDAC, especially when paired with ICB therapy. However, GVAX and related approaches are limited by the use of allogeneic cell lines that do not fully match an individual patient’s tumor antigens and the need to genetically modify these cells to express relevant cytokines. An ideal PDAC vaccine would instead use a patient’s own tumor cells to ensure that all relevant antigens are present and achieve robust immune activation without the need for genetic engineering.

We hypothesized that triggering immunogenic cell death (ICD)[12] of tumor cells *in vitro* could represent a simple and effective way to create an autologous PDAC vaccine that maximizes exposure to tumor antigens and danger signals *in vivo*[13-15]. In particular, tumor cells undergoing ICD display an ‘eat me’ signal, calreticulin (CRT), on the outer membrane[16-20], as well as release damage-associated molecular patterns (DAMPs), including HMGB1 and ATP[21-24], which stimulate dendritic cell maturation and antigen presentation[25-27]. Together, these signals could theoretically convert dead and dying tumor cells into effective stimulators of anticancer immunity. While traditional cancer therapies, such as anthracycline chemotherapy and radiation, can induce ICD in some contexts[28,29], these are not always feasible or optimal for *ex vivo* vaccine preparation. Therefore, we sought to identify a rapid, non-genetic method for inducing ICD in PDAC cells prior to testing vaccine efficacy by injection into a murine tumor model. We first screened various stress induction methods (UV irradiation, hypoxia, heat shock, and chemical oxidative stress) for the ability to maximize ICD hallmarks in murine Panc02 cell lines[30-32]. These analyses identified low-dose H_2_O_2_ as the optimal inducer of oxidative stress, leading to the upregulation of multiple ICD hallmarks and the MHC-I antigen presentation machinery. We then fixed ICD-induced tumor cells and evaluated their therapeutic effects in a syngeneic Panc02 model. The vaccine was tested both prophylactically and therapeutically, as well as in combination with PD-1 blockade, to assess its potential synergies. Together, our results demonstrate that ICD-enhanced tumor cell vaccines can elicit potent immunity against pancreatic cancer by converting ‘cold’ tumors into ‘hot’ and thereby improve responsiveness to ICB.

## MATERIALS AND METHODS

### Cell Lines and Culture

Murine Panc02 pancreatic adenocarcinoma cells (Cytoin Biosciences) were maintained in DMEM with 10% fetal bovine serum and 1% penicillin-streptomycin at 37°C and 5% CO_2_. Murine DC2.4 dendritic cells were cultured in RPMI-1640 supplemented with 10% FBS. All cell lines were routinely tested to confirm mycoplasma-free status. The cells were passaged at 70-80% confluency.

### Immunogenic Cell Death (ICD) Induction

We tested multiple stress conditions to induce immunogenic cell death (ICD) in Panc02 cells[16,33,34]. For irradiation, cells were exposed to 254 nm UV light for 10, 20, or 30 min (in PBS without a lid). Hypoxia was induced by chemical treatment with CoCl_2_ (100 or 200 µM) for 24 h. Heat shock was applied by incubating the cells at 40°C or 42°C for 30 min, followed by recovery at 37°C. Oxidative stress was induced with hydrogen peroxide (H_2_O_2_) at concentrations of 0.03%, 0.05%, 0.1%, or 0.5% over a period of 2 h. After each treatment, cell viability was assessed by trypan blue exclusion, and cells were observed under a light microscope to confirm that >70% of cells remained intact and viable, indicating non-lytic cell death or cellular stress rather than overt lysis. Based on these screens, 0.05% H_2_O_2_ for 2 h was identified as the optimal condition for inducing ICD hallmarks, while acutely preserving >70% cell viability. Oxidative stress was subsequently used for vaccine preparation and downstream analysis.

### Whole-Cell Vaccine Preparation

Panc02 cells treated with 0.05% H_2_O_2_ for 2 h were washed immediately with PBS and fixed with 4% paraformaldehyde (PFA) for 30 min at room temperature. Fixation was performed to ensure that the cells were no longer viable and to preserve the antigenic structures. After fixation, the cells were washed thoroughly to remove PFA, counted, and resuspended in sterile PBS prior to injection as an ICD-induced whole-cell vaccine. Control vaccine preparations included: (1) untreated tumor cells fixed with PFA and (2) PBS-only vehicle control.

### Animal Models and Vaccination Protocols

Animal experiments were approved by the Institutional Ethics Committees (Animal Care Committee, Brock University, AUP 22-11-04; IACUC, Nanyang Technological University, A19019) and adhered to national guidelines. Male C57BL/6 mice (6-8 weeks old) were used for syngeneic tumor studies using the Panc02 cell line. The mice were monitored for tumor growth for 90 days. Tumor size was measured twice weekly using digital calipers and volumes were calculated as (length × width^2^)/2 for an ellipsoid approximation to establish growth curves. Mice were euthanized if the tumor volume exceeded 2000 mm^3^ or if ulceration occurred in accordance with the humane endpoints. Mice were observed for tumor incidence, progression, or regression, with complete regression defined as the disappearance of palpable tumor and confirmation by autopsy. Two vaccination models were used.

Prophylactic Vaccination Model: Mice (n=6 per group) were vaccinated subcutaneously (s.c.) in the left flank with 1×10^6^ fixed Panc02 cells twice on days -14 and -7 (relative to tumor challenge). Three groups were tested: (1) ICD vaccine, H_2_O_2_-treated Panc02 cells fixed with PFA; (2) dead tumor cell vaccine, Panc02 cells fixed with PFA without prior H_2_O_2_ exposure (non-ICD control); and (3) PBS vehicle control. On day 0, all mice were challenged with 2×10^5^ live Panc02 cells subcutaneously in the right flank (contralateral to the vaccination site).

Therapeutic Vaccination Model: Mice (n=6 per group) were subcutaneously injected into the right flank with 2×10^5^ Panc02 cells. After palpable tumors were established (reaching ∼5-8 mm^3^, ∼7 days post-inoculation), the therapy was initiated. Mice were randomized to receive (1) ICD vaccine alone (H_2_O_2_-treated Panc02 vaccine s.c. in the left flank), (2) ICD vaccine + anti-PD-1, (3) anti-PD-1 alone, or (4) PBS vehicle control. The vaccine was administered subcutaneously on day 7, followed by two booster doses on days 14 and 21. For groups receiving anti-PD-1 checkpoint blockade, mice were injected i.p. with 200 µg anti-mouse PD-1 monoclonal antibody (clone RMP1-14, BioXCell) three days after each vaccination (on days 10, 17, and 24), for a total of three cycles. The control groups received an IgG isotype control.

### Immune Response Assays

One week after the final vaccination, blood was collected from the saphenous vein for peripheral blood mononuclear cell (PBMC) isolation. To assess antigen-specific T cell responses, interferon-gamma (IFN-γ) ELISpot assay was performed. Briefly, murine DC2.4 dendritic cells were pulsed with Panc02 whole-cell lysate for 24 h. Isolated PBMCs were then co-cultured with antigen-pulsed DC2.4 cells. The control wells contained PBMCs stimulated with concanavalin A (positive control) or medium alone (negative control). IFN-γ production was detected using a commercial mouse IFN-γ ELISpot kit ( #3321-4HPT-2; Mabtech) according to the manufacturer’s protocol. Spot-forming units (SFU) were enumerated, and the results were expressed as SFU per 10^6^ cells.

### Tumor Immunohistochemistry (IHC)

At the experimental endpoint, the tumors were excised and fixed in 4% paraformaldehyde. Cryosections were subjected to immunohistochemical staining to quantify tumor-infiltrating lymphocytes and natural killer (NK) cells. Briefly, sections were subjected to antigen retrieval, followed by incubation with primary antibodies specific for CD4 (helper T cells), CD8α (cytotoxic T cells), and NCR1 (NKp46, a marker for NK cells). Antibody binding was detected using the VECTASTAIN Elite ABC-Peroxidase Kit with the appropriate secondary antibodies and DAB substrate. The sections were counterstained with hematoxylin. For quantification, five-ten random high-power fields (HPFs) were analyzed per tumor section. The numbers of CD4+, CD8+, and NCR1+ cells were counted in each field, and the average counts per HPF were compared across experimental groups.

### Western Blot and DAMP Release Assays

Panc02 cells treated with or without H_2_O_2_ were analyzed by western blotting to confirm the expression of ICD hallmarks *in vitro*. Total cellular proteins were extracted in RIPA buffer. Equal amounts of protein were separated by SDS-PAGE and then transferred to PVDF membranes. Blots were probed for calreticulin (CRT), ERp57 (an ER chaperone that complexes with CRT), and HMGB1. GAPDH was used as the loading control. To assess HMGB1 release, culture supernatants (conditioned media) were collected after treatment, concentrated, and analyzed by western blotting. The membrane was probed with an anti-HMGB1 antibody, and a positive band in supernatants with concurrent loss of the intracellular HMGB1 band was identified as active release. Band intensities were measured using ImageJ and normalized to the loading control for comparison between the treated and control cells. The antibodies used in the experiments are listed in **Table S1**.

### Immunofluorescence

Surface exposure of calreticulin (ecto-CRT) was evaluated using immunofluorescence imaging. Cells were grown on slides, fixed, stained with anti-calreticulin primary antibody and fluorescent secondary antibodies, and counterstained with DAPI to label the nuclei. Cell surface MHC-I expression was examined by fluorescence microscopy. Treated and control cells on coverslips were stained for H-2K^b^ (green) and nuclei (DAPI, blue) to assess MHC-I expression, respectively.

### Gene Expression Analysis

RNA was extracted from control and H_2_O_2_-treated Panc02 cells using the RNeasy Plus Kit (Qiagen), purified, and subjected to RNA sequencing (RNA-seq). Poly(A)+ RNA libraries were prepared and sequenced on an Illumina NovaSeq X Plus platform (Novogene Corporation Inc., Sacramento, CA, USA) to generate paired-end 150 bp reads. Quality control was performed using FastQC, and the adapter sequences were trimmed using Trimmomatic. Clean reads were mapped to the mouse reference genome (GRCm38) using STAR aligner. Gene-level quantification was performed using featureCounts, and differential gene expression analysis was conducted using DESeq2 to identify genes that were significantly up-or downregulated (adjusted p<0.05, fold-change >2). Gene set enrichment and pathway analyses (KEGG pathways, GO biological processes) were performed on the differentially expressed genes to identify enriched immune-related pathways. Quantitative RT-PCR (qRT-PCR) was conducted to validate the transcriptional changes in key genes. cDNA was synthesized and qPCR was performed using SYBR Green chemistry. Primers targeted ICD-associated genes (e.g. Calreticulin, Erp57/Grp58, Hspa1a/Hsp70, Hmgb1), antigen processing/presentation genes (H2-K1 [MHC-I heavy chain] and Tap2). Gene expression was normalized to that of the housekeeping gene GAPDH, and fold changes were calculated using the 2^-ΔΔCt^ method.

### Statistical Analysis

Data were analyzed using GraphPad Prism 9.3. Normality was checked using the D’Agostino-Pearson or Shapiro-Wilk tests. For two-group comparisons, Student’s t-test was used if the data were normally distributed; otherwise, the Mann-Whitney U test was applied. For multi-group comparisons, one-way ANOVA followed by Tukey’s multiple comparison test was used for parametric data. The Kruskal-Wallis test with Dunn’s post-hoc test was used for non-parametric data. Tumor growth curves were compared using two-way repeated-measures measures ANOVA. Survival curves were plotted using the Kaplan-Meier method and compared using the log-rank test. ELISpot and cell count data were analyzed using one-way analysis of variance (ANOVA). All quantitative data are presented as mean ± standard error of the mean (SEM) unless otherwise stated. Statistical significance was set at P < 0.05.

## Data Availability

The gene expression data generated in this study have been deposited in the Gene Expression Omnibus (GEO) database under accession number **GSE316213**.

## RESULTS

### Oxidative stress induces immunogenic cell death and MHC-I upregulation in Panc02 cells

First, we determined the optimal conditions for inducing immunogenic cell death (ICD) in pancreatic cancer cells. After testing for UV, heat shock, hypoxia (not shown), and oxidative stress, exposure to low-dose hydrogen peroxide (0.05% H_2_O_2_ for 2 h) consistently led to the exposure of multiple ICD hallmarks in murine Panc02 cells without causing significant cell death (Figure 1). Western blot analysis confirmed that H_2_O_2_ treatment led to endoplasmic reticulum (ER) stress and DAMP release, including marked increases in calreticulin (CRT) protein levels (Figure 1A) and upregulation of the ER chaperone ERp57 (Figure 1B) relative to untreated control cells. Furthermore, H_2_O_2_-induced stress led to the active release of the nuclear damage-associated ICD hallmark HMGB1, with protein levels in the intracellular compartment dropping significantly (Figure 1C), whereas concentrations in the culture supernatant were markedly increased (Figure 1D). Since ICD involves the translocation of CRT to the plasma membrane, we also used immunofluorescence to visualize the cellular localization of this protein. Untreated cells displayed low CRT expression, which was confined to the cytosol, whereas H_2_O_2_-treated cells displayed intense staining on the cell surface (ecto-CRT; Figure 1E). Quantification of surface-exposed CRT (mean fluorescence intensity across >150 regions of interest per condition) confirmed a robust increase in ecto-CRT following H_2_O_2_ treatment. Importantly, cells exposed to low-dose H_2_O_2_ did not lyse immediately and maintained integrity throughout the treatment period, allowing efficient and sustained exposure to immunostimulatory signals (rather than uncontrolled release of cellular contents via necrosis). Indeed, when analyzed by qRT-PCR, H_2_O_2_-treated Panc02 cells also displayed upregulation of transcripts for ICD hallmarks CRT, Erp57, HMGB1, and HSP90B1 (fold-changes of ∼1.5-2.5, p<0.05, each; Figure 1G). Together, these data indicate that oxidative stress enhances the tumor cell expression of ICD-related genes and induces a DAMP profile conducive to immune activation, which could potentially act as a cell-based vaccine for PDAC.

**Figure 1.**
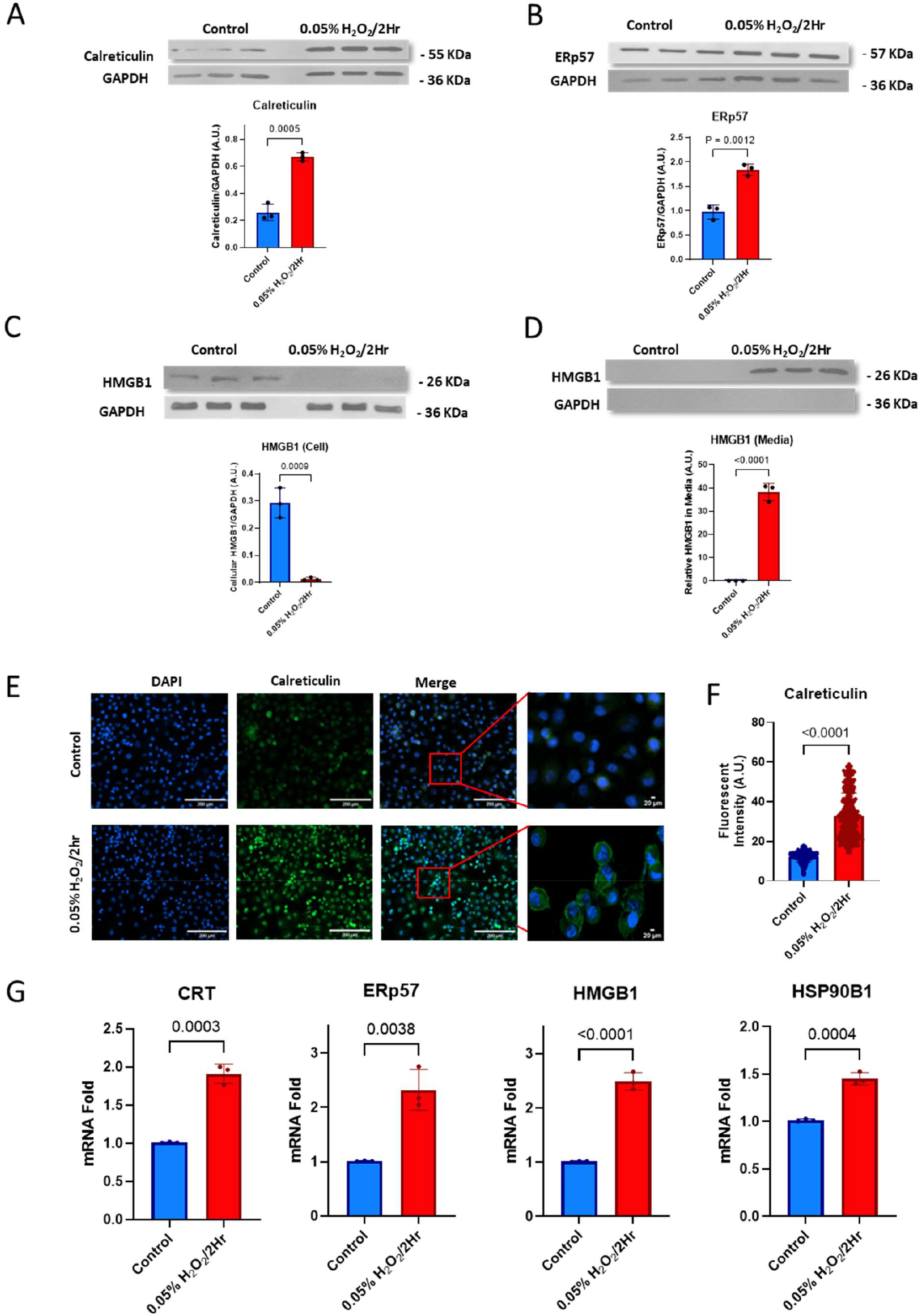
Low-dose H_2_O_2_ induces immunogenic cell death (ICD) in Panc02 cells. Panc02 cells were either exposed to 0.05% H_2_O_2_ for 2 h or left untreated (Control). **(A)** Western blot confirmed a significant increase in total cellular calreticulin in H_2_O_2_-treated cells vs. control. **(B)** Western blot analysis of chaperone protein ERp57 demonstrated a marked increase after H_2_O_2_ treatment, consistent with induction of ER stress. Western blot analysis of HMGB1 in H_2_O_2_-treated cells revealed that protein levels were markedly reduced in the intracellular compartment (**C**), and correspondingly increased in the culture medium (**D**), confirming active HMGB1 secretion/release following treatment. **(E)** Immunofluorescence images of Panc02 cells stained for calreticulin (green) and nuclei (blue). H_2_O_2_-treated cells displayed intense calreticulin staining at the cell surface (ecto-CRT exposure). **(F)** Quantification of calreticulin by Image J analysis (mean fluorescence intensity across >150 regions of interest per condition) confirmed a robust increase in ecto-CRT in H_2_O_2_-treated cells (p<0.0001). **(G)** qRT-PCR analysis of ICD-related genes (CRT, ERp57, HMGB1, HSP90B1) demonstrated significant transcriptional upregulation in H_2_O_2_-treated Panc02 cells compared to controls. These results confirm that low-dose oxidative stress triggers classical ICD markers in pancreatic cancer cells, including DAMPs exposure/release, while preserving cell structure for vaccine use. Data are presented as mean ± SEM. Statistical significance was determined by unpaired Student’s t-test, with P values indicated. n = 3 per condition.

### Oxidative stress reprograms the tumor cell transcriptome into an immunostimulatory state

To determine how oxidative stress reprograms the immunogenic landscape of Pnac02 cancer cells, we performed RNA sequencing of Panc02 cells after treatment with low-dose H_2_O_2_ (0.05% for 2 h). This analysis revealed extensive transcriptional remodeling, with differential expression analysis identifying 1,518 significantly upregulated and 1,844 downregulated genes (padj < 0.05, |log_2_FC| ≥ 1) compared to untreated controls (Fig. 2A). KEGG pathway enrichment analysis of the upregulated genes indicated that the most significantly altered pathways were predominantly metabolic, involving glycolysis/gluconeogenesis, carbon metabolism, and amino acid biosynthesis (Fig. 2B), likely reflecting heightened energetic demands under oxidative stress. Notably, the ‘Influenza A’ pathway was also prominently enriched in H_2_O_2_-treated cells, with a focused heatmap of constituent genes confirming upregulation of multiple antiviral and inflammatory mediators (Fig. 2C). These include key pattern recognition receptors (Ddx58/RIG-I), adaptor proteins (Mavs, Ticam1/TRIF), transcription factors (Irf7, Stat1, Rela/NF-κB), and effector chemokines (Cxcl10, Ccl5), which are characteristic of an antiviral response and are known to generate potent danger signals for immune activation. Accordingly, interrogation of a predefined ICD gene set confirmed that H_2_O_2_ triggered a robust immunogenic program (Fig. 2D). We also observed strong induction of DAMPs, including heat shock proteins (Hspa1a, Hspa1b, Hsp90ab1) and calreticulin (Calr), along with the pro-inflammatory cytokines TNF and the T-cell chemoattractants Cxcl10 and Ccl5. Concurrent upregulation of Irf7, Stat1, and the autophagy-related gene Atg7 further supported that low-dose H_2_O_2_ can induce a comprehensive ICD response in Panc02 cells. Finally, we assessed the transcriptional regulation of the MHC-I antigen presentation pathway in response to H_2_O_2_ treatment, which significantly increased the expression of subunits H2-T23, H2-K1, and B2m, and essential chaperones and components of the peptide-loading complex (Canx, Hspa1a, Hspa1b, and Hsp90ab1) (Fig. 2E). This coordinated upregulation indicated an enhanced cellular capacity for antigen processing and stable MHC-I surface expression. Collectively, our transcriptomic data demonstrated that sublethal oxidative stress reprograms Panc02 cells into a potent immunogenic state, characterized by the induction of an antiviral danger program, canonical ICD signature, and enhanced antigen presentation machinery.

**Figure 2.**
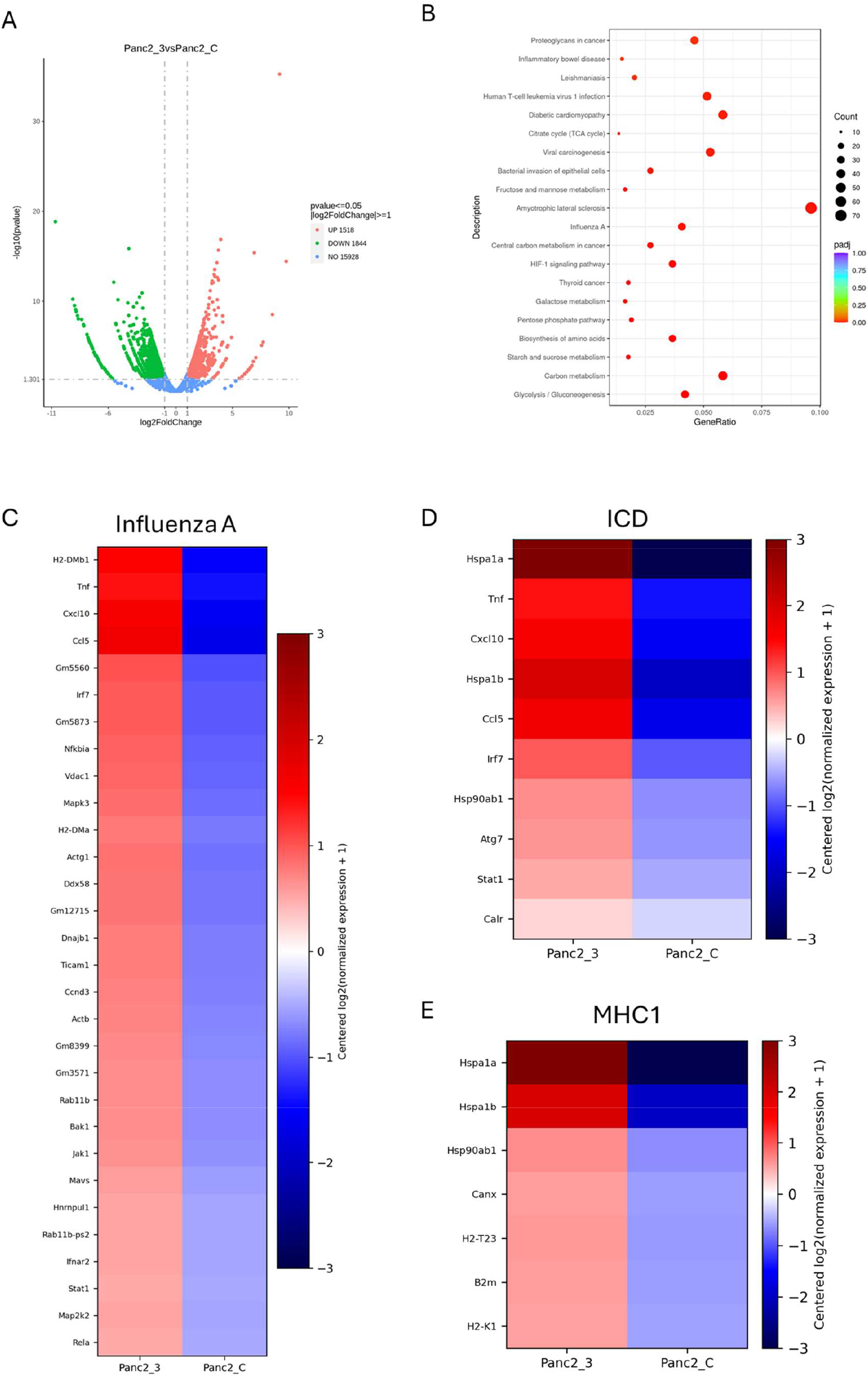
Oxidative stress reprograms the Panc02 transcriptome towards immunogenicity. **(A)** Volcano plot of differentially expressed genes in H_2_O_2_-treated Panc02 cells (Panc2_3) versus untreated controls (Panc2_C). Horizontal dashed line denotes padj = 0.05; vertical dashed lines indicate log_2_ fold-change (log_2_FC) ±1. Red dots represent significantly upregulated genes (n = 1,518), green dots show significantly downregulated genes (n = 1,844), and blue dots indicate non-significant genes (n = 15,928). (B) KEGG pathway dot-plot of significantly enriched pathways among the upregulated genes (padj < 0.05). Dot size reflects the number of genes in each pathway, and dot color indicates adjusted p-value. The most enriched pathways were metabolic (e.g. glycolysis/gluconeogenesis, carbon metabolism), with Influenza A as a representative immune / antiviral pathway. (C) Heat-map of upregulated genes in the ‘Influenza A’ pathway. Coordinated upregulation of key antiviral and inflammatory mediators, including pattern recognition receptors (Ddx58/RIG-I), adaptor proteins (Mavs, Ticam1/TRIF), transcription factors (Irf7, Stat1, Rela/NF-kappaB), and effector chemokines (Cxcl10, Ccl5), characteristic of an antiviral ‘danger’ program. (D) Heat-map of the immunogenic cell death (ICD) gene panel. A robust ICD signature was evidenced by strong induction of DAMPs including Heat Shock Proteins (Hspa1a, Hspa1b, Hsp90ab1), Calreticulin (Calr), pro-inflammatory cytokine TNF, T-cell chemoattractants Cxcl10 and Ccl5, together with upregulation of Irf7, Stat1, and Atg7. (E) Heat-map of MHC-I antigen presentation pathway genes. Enhanced expression of the MHC-I subunits (H2-T23, H2-K1), light chain B2m, and essential chaperones/components of the peptide-loading complex (Canx, Hspa1a, Hspa1b, Hsp90ab1) consistent with increased capacity for antigen processing and stable MHC-I surface expression. All heat-maps display centred log_2_ (normalized expression + 1) values.

### H_2_O_2_ treatment promotes tumor cell antigen presentation

Immunofluorescence analysis of MHC-I expression in Panc02 cells confirmed these transcriptomic findings. As shown in Figure 3A, untreated Panc02 cells displayed modest MHC-I (H-2K^b^) expression, whereas H_2_O_2_-treated cells showed intense MHC-I surface staining, suggesting the distribution of peptide-MHC complexes to the plasma membrane following stress. Quantification across multiple images confirmed a significant increase (∼3-fold) in the mean MHC-I fluorescence intensity per cell in the H_2_O_2_ group compared to that in the control (Figure 3B). Moreover, qRT-PCR of H2-K1 and the antigen-processing gene Tap2 demonstrated coordinated upregulation (Figure 3C), indicating that oxidative stress not only induces DAMP release but also upregulates the antigen presentation machinery. Collectively, these data suggest that H_2_O_2_-treated tumor cells are primed for immunogenicity, including higher levels of DAMPs and surface MHC-I expression, to recruit and activate antigen-presenting cells and cytotoxic CD8 + T cells. Having established the immunogenic properties of H_2_O_2_-treated Panc02 cells *in vitro*, we tested their efficacy as a vaccine *in vivo*.

**Figure 3.**
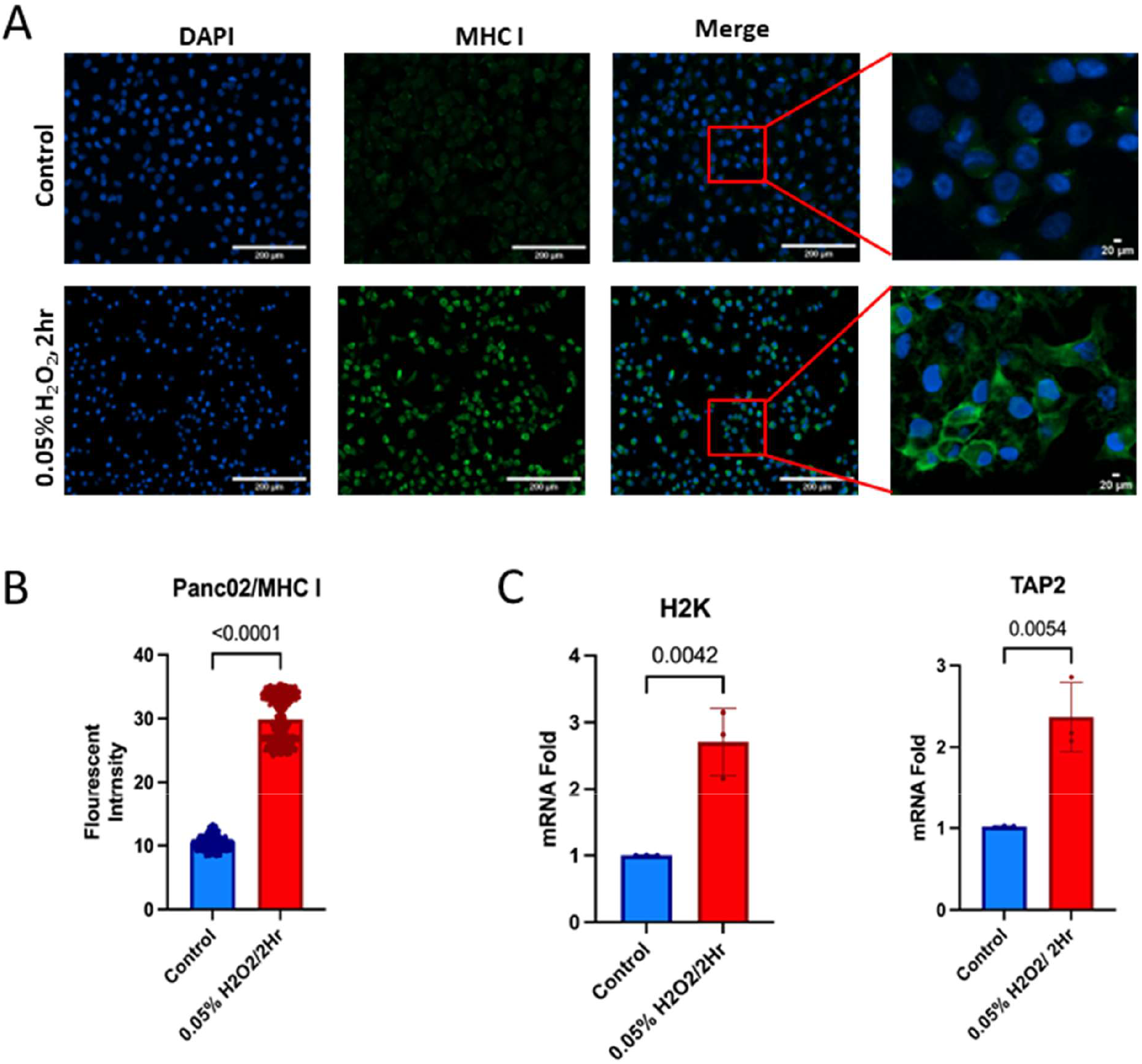
Oxidative stress upregulates antigen presentation machinery in Panc02 cells. (A) Immunofluorescence staining of surface MHC-I (H-2K^b^, green) on Panc02 cells. Upper: untreated control cells show faint surface MHC-I. Lower: 0.05% H_2_O_2_-treated cells display bright MHC-I staining at the plasma membrane. Nuclei are counterstained with DAPI (blue). (B) Quantification of MHC-I fluorescence intensity on the cell surface from multiple images (150 ROI per image, n=3 independent experiments). H_2_O_2_-treated cells display ∼3-fold higher intensity of MHC-I relative to untreated cells (p<0.0001), indicating enhanced antigen presentation potential. (C) Expression levels of antigen presentation genes as measured by qRT-PCR relative to control. H-2K^b^ (MHC-I heavy chain) transcript was increased ∼3-fold in H_2_O_2_-treated cells. Tap2 (peptide transporter) was also significantly upregulated (∼2.5-fold, p=0.0054). Data shown are mean values ± SEM (p values by Mann-Whitney U test), n=3.

### Prophylactic vaccination with ICD-induced PDAC cells elicits protective immunity *in vivo*

To evaluate whether the ICD-enhanced PDAC vaccine could protect against tumor challenge *in vivo*, we pretreated mice with two injections of H_2_O_2_-treated and fixed Panc02 cells (ICD vaccine), control fixed cells only (dead tumor cells without ICD), or a PBS vehicle control prior to challenge with live Panc02 cells (Figure 4A). Mice vaccinated with the ICD-treated whole-cell vaccine exhibited a marked delay in tumor onset and sustained control of tumor growth compared to both control groups (Figure 4B). By day 40 post-challenge, mice in the PBS and dead cell control groups developed large, palpable tumors (mean volume of approximately 100 mm^3^), whereas mice receiving the ICD vaccine displayed only small nodules or no detectable tumors. Consistent with improved tumor control, Kaplan-Meier survival analysis demonstrated a significant survival advantage in the ICD-vaccinated group relative to the controls (log-rank test; Figure 4C). Five of the six mice vaccinated with ICD-treated cells survived throughout the 90-day observation period, whereas all mice in the PBS and dead cell control groups reached humane endpoints by day 60 and day 70, respectively. Vaccination with fixed dead tumor cells lacking prior H_2_O_2_-induced ICD resulted in only a modest delay in tumor progression compared to PBS controls, and all mice in this group ultimately developed tumors and succumbed to disease before day 70. Together, these findings indicate that administration of non-immunogenic killed Panc02 cells provides minimal protection, and that prior induction of ICD by H_2_O_2_ treatment is critical for effective prophylactic antitumor immunity.

**Figure 4.**
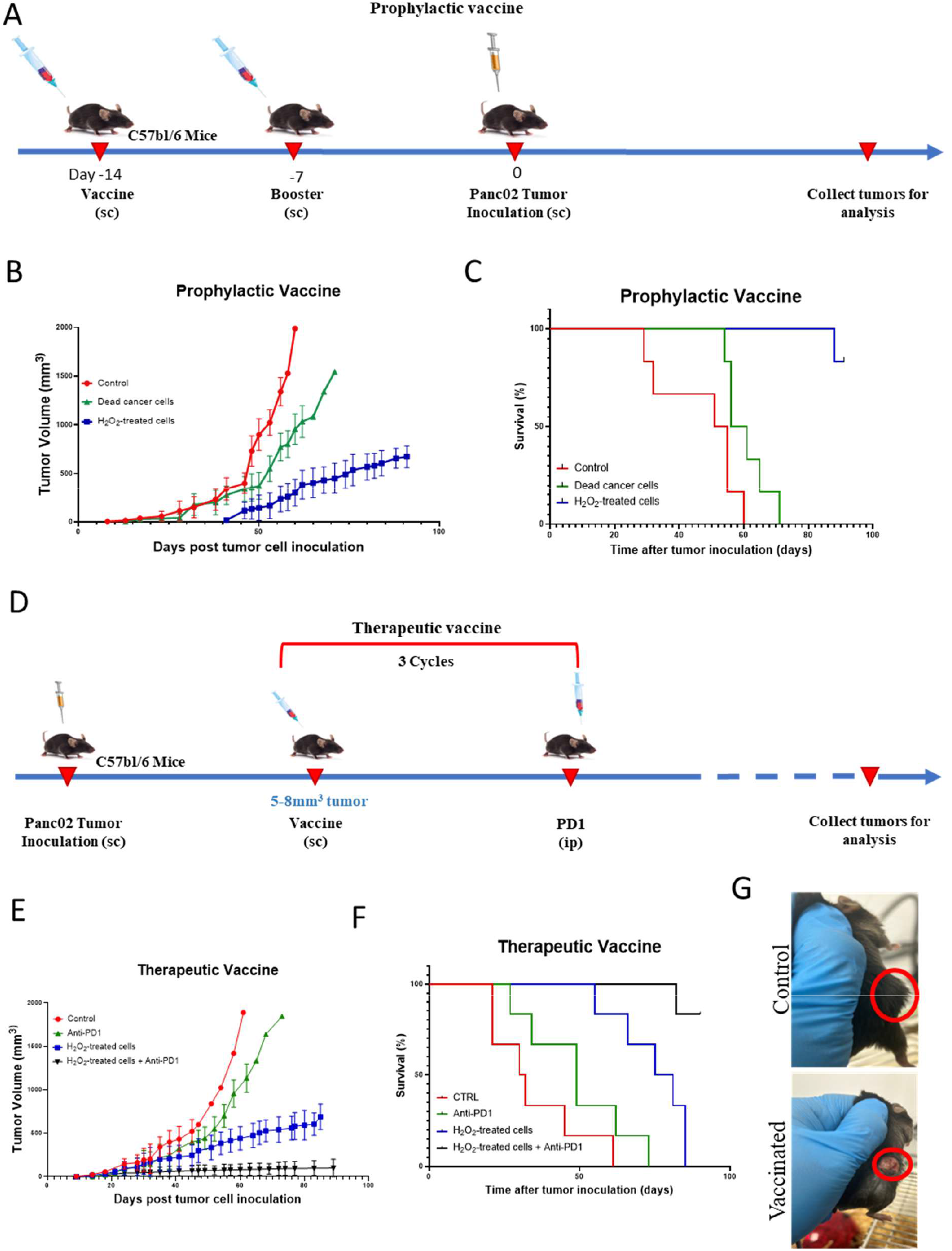
H_2_O_2_-treated Panc02 cell vaccine elicits robust anti-tumor immunity *in vivo*. **(A)** Schematic of the prophylactic vaccination protocol: Mice were vaccinated subcutaneously (s.c.) in the left flank on days -14 and -7 with H_2_O_2_-treated and fixed Panc02 cells (ICD vaccine), or control dead cells, or PBS alone. Tumors were established by subsequent challenge with live Panc02 cells injected in the right flank on day 0. **(B)** Tumor growth curves in the prophylactic model. Mice receiving the ICD vaccine (blue) were largely protected from rapid tumor growth, with 5 of 6 mice remaining alive throughout the experiment (90 days). Vaccinated mice showed significantly delayed growth compared to control groups (non-ICD dead cells, green; PBS, red). Data represent mean tumor volume ± SEM. **(C)** Survival in the prophylactic models. Survival differences were assessed using the log-rank (Mantel-Cox) test. P values for pairwise comparisons: Control vs. dead cancer cells, p = 0.0252; Control vs. H_2_O_2_-treated cancer cells, p = 0.0006; dead cancer cells vs. H_2_O_2_-treated cancer cells, p = 0.0005. **(D)** Schematic of the therapeutic vaccination protocol. Mice with established Panc02 tumors on the right flank received ICD vaccine s.c. in the left flank, either alone or in combination with anti-PD-1 antibody. **(E)** Tumor growth curves in the therapeutic setting. The ICD vaccine monotherapy (blue) slowed tumor progression compared to controls. Combining ICD vaccine with anti-PD-1 (black) induced tumor regression or slow tumour growth, significantly improved tumour control compared to vaccine alone. Anti-PD-1 monotherapy (green line) had minimal effect on tumor growth and was only slightly delayed compared to the PBS vehicle-only control. Data shown are mean values ± SEM. **(F)** Survival curves of tumor-bearing mice. Combining ICD vaccine with anti-PD-1 significantly extended survival, with 5 out of 6 mice remaining alive at the day 90 endpoint. Survival differences were assessed using the log-rank (Mantel-Cox) test. P values for pairwise comparisons: Control vs. anti-PD-1, p = 0.091; Control vs. H_2_O_2_-treated cancer cells, p = 0.0015; Control vs. H_2_O_2_-treated cancer cells + anti-PD-1, p = 0.0005; anti-PD-1 vs. H_2_O_2_-treated cancer cells + anti-PD-1, p = 0.0005. **(G)** Representative photographs of the skin over the tumor site. A localized inflammatory reaction (red circle) was observed at the tumor site (right flank) 3 days post-vaccination with H_2_O_2_-treated cells administered via the left flank. This reaction was absent in control-vaccinated mice. Data are representative of n=6 mice per group. Statistical significance was determined by log-rank test.

### Therapeutic vaccination slows tumor growth and synergizes with PD-1 blockade

Next, we tested the PDAC cell vaccine in a therapeutic setting and modeled the treatment of established pancreatic tumors. Mice bearing palpable Panc02 tumors (∼5-8 mm^3^ at treatment initiation, approximately 7 days after tumor cell inoculation) were randomized to receive the ICD vaccine alone or in combination with anti-PD-1 antibody, anti-PD-1 antibody alone, or corresponding control treatments (Figure 4D). As expected, the untreated tumors grew aggressively, reaching the humane endpoint by approximately 5-6 weeks (Figure 4E, red curve). Mice treated with H_2_O_2_-treated ICD vaccine (Figure 4E, blue curve) displayed significantly slower tumor growth relative to controls, with ∼50% lower tumor volume compared to PBS vehicle by day 40. However, the ICD cell vaccine alone was insufficient to induce tumor regression, and anti-PD-1 treatment alone (green curve in Figure 4E) resulted in only a modest delay in tumor growth compared with the PBS vehicle control (red curve), consistent with clinical observations that PD-1 blockade alone is ineffective in PDAC. The combination of the ICD vaccine and anti-PD-1 therapy resulted in superior therapeutic efficacy. By day 60, the average tumor volume in the combination therapy group grew slowly and remained palpable, whereas tumors in the vaccine only group were >100 mm^3^ when control tumors reached experimental or humane endpoints (growing rapidly to 2000 mm^3^). Figure 4E shows that combination therapy (black curve) led to very slow tumor growth and small tumor volume after ICD vaccine and anti-PD-1 treatment, whereas vaccine alone (blue curve) only stabilized or slowed growth. This synergy translated into improved overall survival (Figure 4F), with mice receiving both vaccine and PD-1 blockade achieving 5 of 6 mice survival at 90 days, compared with a median ∼78 days survival among animals receiving vaccine alone, ∼49 days for anti-PD-1 blockade alone, and ∼38 days for control mice. These results demonstrate that while ICD-based vaccines can significantly prime immune responses *in vivo*, additional checkpoint inhibition is required to fully realize anti-tumor immune responses, leading to effective treatment in many individuals.

Another observation was localized skin inflammation at the site of the tumor mass in ICD-vaccinated mice. Approximately 3 days after vaccination, mice that received H_2_O_2_-treated cells developed redness above the tumor mass, which did not occur in mice injected with PBS or non-ICD dead cells. Figure 4G shows a representative image of erythema and induration visible on the right flank (tumor site), which we interpret as a positive sign of local anti-tumor responses [35-37]. We also monitored the mice for adverse effects throughout the treatment period. In addition to the transient inflammation noted above, the vaccine was well tolerated. Mice receiving the combination therapy did not exhibit obvious additional toxicity, suggesting that the injected ICD vaccine did not trigger harmful autoimmunity.

### ICD vaccine induces systemic tumor-specific T cell responses

To confirm that the PDAC vaccine was generating tumor-specific T cell responses, we performed IFN-γ ELISpot assays using peripheral blood mononuclear cells (PBMCs) from treated mice. Compared to control mice, animals vaccinated with H_2_O_2_-treated Panc02 cells displayed a substantially higher frequency of IFN-γ-secreting T cells after re-stimulation with Panc02 antigen-loaded DC2.4 cells (Figure 5). PBMC from ICD-vaccinated mice yielded ∼300 IFN-γ spots per 10^6^ cells, whereas those from PBS-treated-or dead-cell-treated mice produced only 100-150 spot counts (in response to the same tumor antigen stimulus). Combining PD-1 blockade with the vaccine did not significantly increase the number of PBMC producing IFN-γ in three cycles of therapy, indicating that the vaccine alone had already maximized this response, but it did appear to sustain T cell functionality within the tumor (as evidenced by the anti-tumor responses and improved survival, as shown above, and T-cell tumor infiltration data below). These results demonstrate that circulating T cells capable of recognizing Panc02 antigens were present in vaccine groups but were rare in control animals, suggesting that ICD vaccines act systemically to prime T cells in lymphoid organs that subsequently traffic to the tumor site.

**Figure 5.**
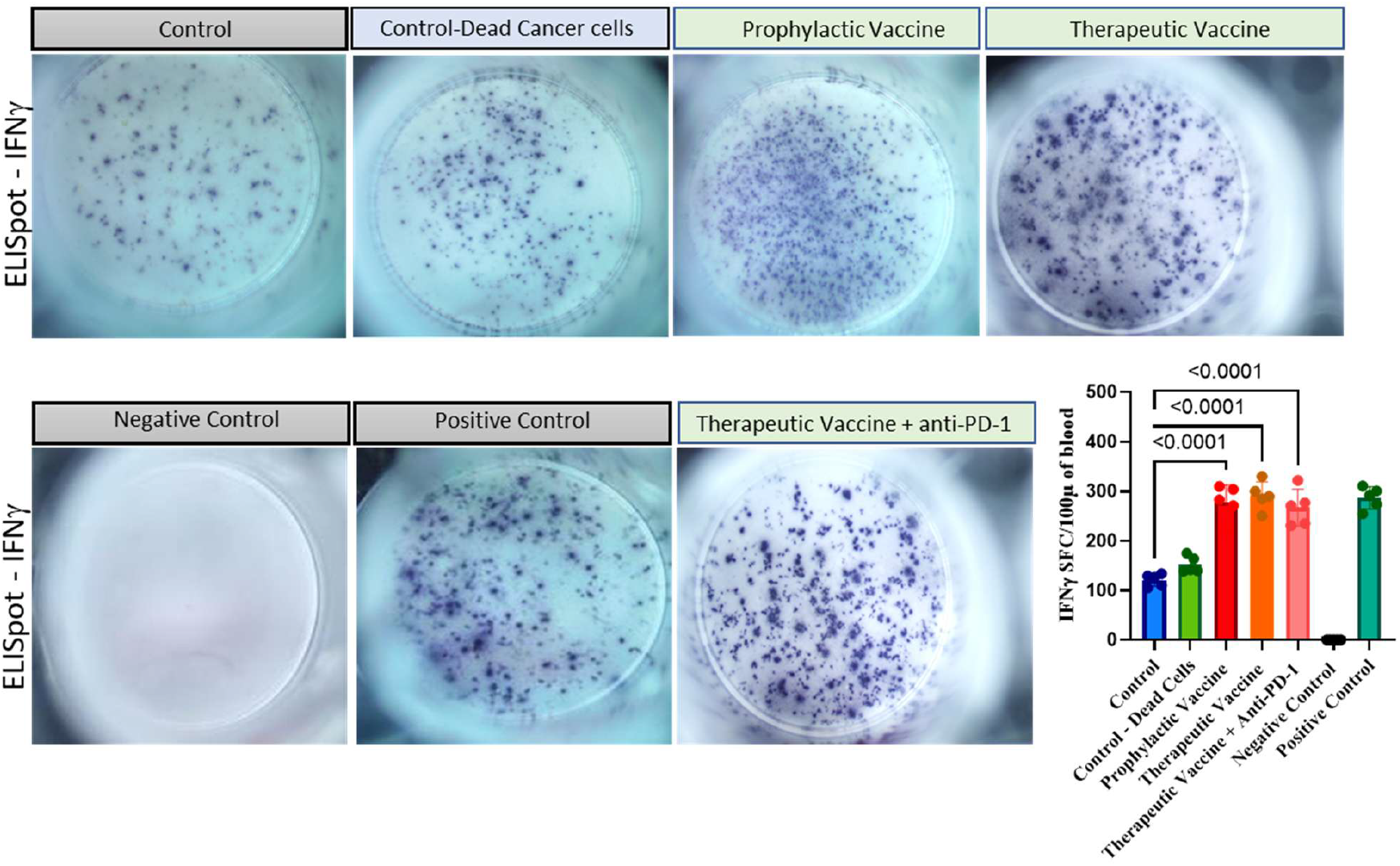
H_2_O_2_-treated PDAC vaccine induces strong tumor-specific IFN-γ T cell responses. Representative IFN-γ ELISpot results from PBMCs of vaccinated mice 3 weeks after therapy initiation. PBMCs from an ICD-vaccinated mouse show numerous spots upon co-culture with DC2.4 cells pulsed with Panc02 tumor lysates, indicating IFN-γ release by antigen-specific T cells. In contrast, PBMCs from a control mouse produced only a few spots. Bar graph: Quantification of ELISpot results (spots per 1×10^6^ cells) for each group (n=5 mice/group). ICD vaccine elicited ∼300 IFN-γ SFU/10^6^ PBMC on average, significantly above the ∼120 SFU detected in controls (p<0.001). Vaccinated mice that also received anti-PD-1 achieved similarly high spot counts, confirming that the vaccine is the main driver of T cell priming. Mice that received dead cells vaccine showed only slight increase in spots, consistent with their weak protective effects. These ELISpot data indicate that the ICD-based vaccine generated a robust T-cells response against tumor antigens *in vivo*.

### ICD vaccination plus PD-1 blockade promotes tumor infiltration by T- and NK-cells

To assess how ICD-based vaccination reshapes the pancreatic tumor microenvironment, we next performed immunohistochemistry on tumor sections from the five treatment groups (PBS, dead cells only, prophylactic ICD vaccine, therapeutic ICD vaccine, and therapeutic ICD vaccine + anti-PD-1) (Figure 6). Consistent with the immune-exclusionary profile of untreated Panc02 tumors, PBS controls contained almost no CD4 + or CD8 + T cells and NCR1^+^ NK cells were virtually absent. Similarly, Panc02 tumors from mice receiving the dead-cell vaccine showed minimal lymphocytic infiltration, indicating that antigen exposure alone only slightly increased the effector cells in the TME in the absence of immunogenic cell death. In contrast, ICD vaccination induced a marked increase in CD4 + helper T-cell infiltration. Quantification of the% DAB-positive area showed that CD4 + staining increased from <1% in PBS/dead-cell controls to ∼2% with prophylactic ICD vaccination, ∼5% with therapeutic vaccination, and ∼6% with the therapeutic vaccine combined with PD-1 blockade. In ICD-treated tumors, CD4 + T cells invaded deep into the tumor, with the densest infiltrates observed after combination therapy.

**Figure 6.**
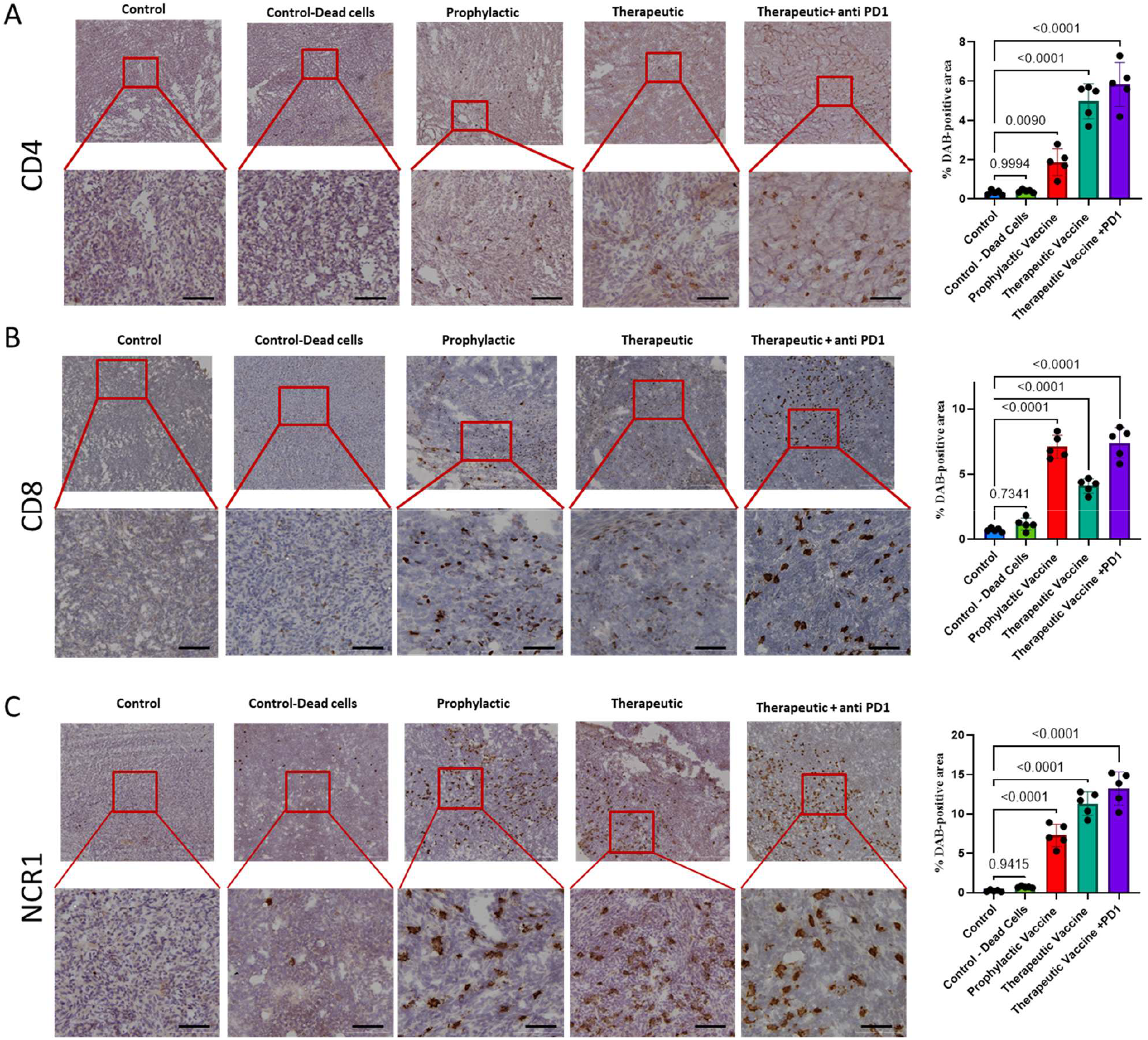
ICD mono/combo therapy increases lymphocyte infiltration of pancreatic tumors. **(A)** Representative CD4^+^ T cell immunohistochemistry of tumor sections from different treatment groups. PBS-treated control tumors contained virtually no CD4^+^ cells. Tumors from a mouse vaccinated with dead (non-ICD) cells displayed only rare CD4^+^ lymphocytes. In contrast, a tumor from ICD-vaccinated mouse exhibited numerous CD4^+^ T cells (brown staining) infiltrating the tumor microenvironment. A tumor from the vaccine + anti-PD-1 combination group exhibited even denser CD4^+^ infiltrate. Insets below each panel show higher magnification of CD4^+^ cell areas (brown spots). Quantification of CD4^+^ T cells using % DAB positive area from Image J. **(B)** Representative CD8^+^ T cell IHC (brown) in tumors. Control and dead-cell tumors essentially lacked CD8^+^ cells. The ICD vaccine tumor contained many infiltrating CD8^+^ T cells, and the combo therapy tumor was heavily infiltrated with CD8^+^ cells. Quantification of CD8^+^ T cells in tissue showing a dramatic increase with ICD vaccine conditions compared to controls. **(C)** Representative NK cell (NCR1^+^) IHC. NK cells (brown) were largely absent from control tumors, but significant numbers were present in prophylactic ICD vaccine TME, and further increased in therapeutic ICD vaccine TME. In the combo treatment tumors, NK cells were more numerous in the TME. Quantification of NCR1^+^ NK cells using Image J % DAB positive area. Each quantification represents data from n = 5 mice per condition and is presented as mean ± SD. Statistical significance was determined by one-way ANOVA followed by Tukey’s post hoc test for all panels. Scale bar: 50 µm.

A similar pattern was observed for CD8 + T cells. Control tumors contained almost no detectable CD8 + staining, whereas prophylactic ICD vaccination increased the CD8 + DAB area to ∼7% compared with ∼1-2% in controls. Therapeutic ICD vaccination produced ∼4% CD8 + area, and the addition of PD-1 blockade resulted in the greatest infiltration (>7.5% DAB-positive area). CD8 + cells commonly appeared within the tumor cores, which is consistent with active cytotoxic responses. NCR1^+^ NK cells followed the same trend; they were essentially absent in PBS and dead-cell controls, but increased markedly following ICD vaccination. Prophylactic ICD vaccination increased NK staining approximately 7-fold over PBS only, therapeutic ICD vaccination increased NK cell density ∼11-fold, and combination therapy with anti-PD-1 further increased NK cell infiltration by ∼13-fold compared to controls (p < 0.0001). The presence of intratumoral NK cells is biologically significant and contributes to antitumor immune responses by promoting inflammatory cytokine production, dendritic cell activation, and cytotoxic T cell immunity[38-40]. We also noted the presence of tertiary lymphoid structures (organized clusters of B and T cells with APCs) in some vaccinated tumors (not shown), consistent with therapy-induced lymphoid neogenesis, as has also been reported for GVAX[41].

In summary, the results from our tumor models confirmed that ICD-induced vaccines can overcome the immune-exclusion features of PDAC tumors. By providing both a source of tumor antigens and endogenous adjuvants (DAMPs), the vaccine enabled both T and NK cells to penetrate the tumor. However, therapeutic efficacy was limited until combined with checkpoint blockade, which enabled potent anti-tumor responses and even complete regression in some treated mice. Thus, our data provide preclinical proof of concept that oxidative stress-induced ICD-based autologous vaccines can synergize with ICB to effectively treat pancreatic cancer.

## DISCUSSION

Pancreatic cancer is a prototypical ‘cold’ tumor possessing numerous suppressive features that result in poor immunotherapy outcomes[42]. This study reports a novel method for enhancing pancreatic tumor cell immunogenicity by exposure to oxidative stress and subsequent ICD. Unlike earlier PDAC vaccine approaches that require genetic engineering (e.g., GVAX’s GM-CSF expression), our method uses a simple chemical stressor to create a whole-cell vaccine with potent adjuvant effects *in vivo*. We observed that 2 h of exposure to low-dose H_2_O_2_ was sufficient to trigger ICD hallmarks in Panc02 cells, including upregulation of ER chaperones (ecto-CRT, ERp57) and rapid release of HMGB1, likely to recruit and activate antigen-presenting cells *in vivo*. Accordingly, our transcriptomic data confirmed the induction of a canonical ICD signature, including the upregulation of heat-shock protein genes (Hspa1a/b, Hsp90), chemokines (Cxcl10, Ccl5), and TNF (Figure 2D). Interferon-related genes (Irf7, Stat1) and NF-κB regulators are also elevated to create an ‘antiviral’ inflammatory state, which is increasingly recognized to exert a critical influence on tumor progression and treatment outcomes[43,44].

In parallel with DAMP release, oxidative stress also increased the surface levels of MHC-I in Panc02 cells, and RNA-seq/qPCR analyses confirmed the higher expression of H2-K1 and Tap2. By increasing MHC-I activity in tumor cells, oxidative stress likely reveals more tumor antigen peptides for immune cell engagement. Accordingly, vaccination with H_2_O_2_-treated cells provided robust protection and therapeutic efficacy in murine models *in vivo*. Prophylactically, two doses inhibited tumor establishment or significantly delayed outgrowth in the syngeneic model, whereas a control vaccine of fixed, non-immunogenic cells produced only minimal protection, characterized by a modest delay in tumor growth (Figure 4B-C). Therapeutically, the ICD vaccine slowed tumor progression (Figure 4E), but the impact was markedly enhanced when combined with PD-1 blockade, leading to tumor regression and dramatically extending survival in a significant proportion of mice (Figure 4F). This synergy with checkpoint inhibitors resembles that reported for other vaccine strategies, for example, GVAX, suggesting that both CD8 + T cell priming and release from PD-1 inhibition are required to achieve maximum anti-tumor responses. Indeed, our ELISpot assays revealed a ∼2-3-fold expansion of tumor-specific IFNγ responses by CD8 + T cells from vaccinated mice (Figure 5). IHC analysis of tumors also confirmed that vaccination triggered a dramatic change in the immune landscape, transitioning from an ‘immune desert’ to an ‘inflamed’ profile. H_2_O_2_-vaccinated tumors displayed extensive infiltration of both CD4 + helper T cells and CD8 + cytotoxic T cells (Figure 6A-B), as well as NCR1^+^ NK cells (Figure 6C), indicating broad activation of anti-tumor immunity.

Pancreatic cancer is known to harbor a highly immunosuppressive tumor microenvironment, in which PD-1/PD-L1 signaling limits T-cell effector function. Consistent with this established mechanism, the combination of an ICD vaccine with anti-PD-1 markedly enhanced therapeutic efficacy. These findings suggest that PD-1 blockade is required to relieve checkpoint-mediated suppression and enable vaccine-primed T cells to exert effective antitumor activity in pancreatic tumors[45]. Using whole cells, the vaccine can present many available tumor antigens, including mutant neoantigens and potentially non-mutant tumor-specific epitopes arising from aberrant processing. Recent studies have highlighted that cancer cells generate neoantigens not only from DNA mutations, but also via abnormal RNA splicing, cryptic translation, and post-translational modifications[46-49], which would be missed by mutation-centric vaccines. Our RNA-seq data also confirmed the upregulation of antigen-processing genes, implying that peptides from these diverse antigens could be efficiently presented. Compared to peptide or RNA vaccines, which target only a few predicted epitopes, this whole-cell strategy can broaden the repertoire of tumor antigens and may elicit T cell responses against multiple targets.

Compared with previous PDAC vaccine approaches, our H_2_O_2_ conditioning method offers several practical advantages. Traditional GVAX requires gene transfer and culture of secreted allogeneic cancer cell lines, whereas neoantigen vaccines require genome sequencing, epitope prediction, and custom peptide synthesis, both of which are costly and time-consuming. In contrast, an ICD vaccine can be generated on demand from a patient’s own tumor tissue obtained by biopsy or surgical resection, ensuring full autologous MHC compatibility and requiring only brief *ex vivo* chemical induction of immunogenic cell death followed by fixation. By avoiding the need for genetic engineering, this simple protocol can dramatically reduce the delay from the initial tumor biopsy to patient vaccination. Moreover, the potent stimulatory properties of ICD may reduce the need for additional adjuvants. Indeed, we observed that the vaccine alone was sufficient to elicit strong host responses, although exploring possible combinations with TLR agonists would be a logical next step in future work.

A limitation of our study was that we tested the tumor vaccine in a single syngeneic model (Panc02 in C57BL/6 mice). Although this model is widely used in preclinical PDAC immunotherapy research, human tumors are far more heterogeneous. Validation in additional models, including humanized mice or patient-derived xenografts, would therefore strengthen translational relevance. We also used paraformaldehyde to fix vaccine cells for safety reasons. In clinical settings, radiation or other sterilization methods can replace PFA. The specific dose and timing of H_2_O_2_ exposure may also require optimization in different cell types. Finally, we did not measure all possible immune parameters in these proof-of-concept experiments; therefore, a more granular analysis of T-cell epitopes and memory formation using single-cell RNA-seq would be informative. Despite these limitations, our data clearly showed that oxidative stress-induced ICD is a viable strategy for generating an effective whole-cell cancer vaccine. By harnessing the full antigenic complexity of tumor cells, this approach can overcome intrinsic resistance mechanisms in PDAC and potentially form the basis of personalized immunotherapy in humans. For example, autologous tumor tissue collected at biopsy or at the time of tumor resection could be treated *ex vivo* with a defined oxidative stress protocol and then fixed and re-infused as a custom vaccine in combination with checkpoint inhibitors. This strategy can complement the existing cancer immunotherapy pipelines.

## CONCLUSION

We present a simple, scalable approach to convert pancreatic tumor cells into a potent vaccine using oxidative stress-induced ICD. Low-dose H_2_O_2_ treatment exposes DAMPs and upregulates MHC-I, rendering Panc02 tumor cells highly immunogenic. Vaccination with these cells primes broad tumor-specific T cell responses and controls tumor growth *in vivo*, particularly when combined with PD-1 blockade. This method avoids the need for complex antigen discovery or genetic modification, thereby harnessing the patient’s own tumor antigen repertoire for personalized immunotherapy. Our findings suggest that chemically induced ICD vaccines could be a promising method for overcoming immune evasion in ‘cold’ tumors.

## Supporting information

Antibodies used in the experiments

## ACKNOWLEDGEMENTS

We thank the animal facility staff for their excellent care and maintenance of the experimental animals used in this study. This work was supported in part by the Canadian Institutes of Health Research Tier1 Canada Research Chair (CRC-2020-00263), Canadian Institutes of Health Research Project Grant (PJT-186091), Natural Sciences and Engineering Research Council of Canada Discovery Grant (RGPIN-2023-04304), Canada Foundation for Innovation Grant (41454 and 44115), Ontario Research Fund, start-up research grant from Brock University, and Singapore National Medical Research Council (NMRC/OFIRG/0003/2016).

