## Supplementary material for "Oxidative Stress-Induced Immunogenic Cell Death Enhances Whole-Cell Vaccine Efficacy in a Syngeneic Pancreatic Cancer Model": Antibodies used in the experiments

**Table S1. Antibodies used in this study**

| **Application** | **Marker/Target** | **Manufacturer/Vendor** | **Catalog No.** | **Clone** |
| --- | --- | --- | --- | --- |
| IF | MHC I* | In-house generated (hybridoma-derived) | N/A | 24.14.85 (anti–H-2Kᵇ/Dᵇ) + Y3 (anti–H-2Kᵇ) |
| IF | Calreticulin | Abcam | ab92516 | EPR3924 |
| IHC | CD4 | Cell Signaling Technology | 25229S | D7D2Z |
| IHC | CD8 | Cell Signaling Technology | 98941S | D4W2Z |
| IHC | NCR1 | Abcam | ab233558 | EPR23097-35 |
| WB | Calreticulin | Abcam | ab92516 | EPR3924 |
| WB | ERp57 | Cell Signaling Technology | 2881S | G117 |
| WB | HMGB1 | Abcam | ab79823 | EPR3507 |
| WB | GAPDH | Abcam | ab8245 | 6C5 |
| Injection | Anti–PD-1 | Bio X Cell | CP151 | RMP-14 |

* Murine MHC class I was detected using an in-house-generated primary antibody consisting of a mixture of monoclonal antibodies 24.14.85 (anti–H-2Kᵇ/Dᵇ) and Y3 (anti–H-2Kᵇ).
